# Temperature signals drive grass secondary cell wall thickening

**DOI:** 10.1101/2025.04.03.647122

**Authors:** Greg A. Gregory, Joshua H. Coomey, Bahman Khahani, Didier Gonze, Serene Omran, Kathryn A. McGillivray, Conrad E. Stewart, Kira A. Gardner, David Follette, Samuel P. Hazen

## Abstract

In grasses, stem elongation is driven by intercalary meristems at node-internode junctions, where cells divide, elongate, and in some cell types secondary wall maturation. Cellulose is the predominant polymer in plant cells and the most abundant biopolymer on Earth. It is synthesized at the plasma membrane by multi-protein complexes that include CELLULOSE SYNTHASE A (CESA) proteins. To investigate the spatiotemporal regulation of cellulose deposition during development, we developed a *CESA8* luciferase gene expression reporter system in *Brachypodium distachyon*. High bioluminescence was observed in stem nodes, a specific region of elongating internodes, and the inflorescence, indicating sites of active secondary wall deposition. Within internodes, luminescence followed a distinct pattern, with a "dark zone" directly above the node with minimal signal, followed by a "bright zone" approximately 5 mm above the node where bioluminescence peaked. Histological, biophysical, and transcript analysis confirmed that luminescence intensity correlates with thickened secondary cell walls, increased cellulose crystallinity, and elevated *CESA8* transcript levels. Time-lapse imaging revealed that *CESA8* expression follows a robust diurnal rhythm governed by thermocycles alone, with peak expression occurring in the early morning. Temperature pulse experiments revealed an immediate but transient response of *CESA8* to temperature shifts, which we modeled as an incoherent feed-forward loop. Finally, we found a strong correlation between *CESA8* expression and stem elongation, highlighting the role of secondary cell wall thickening in supporting upright growth. These findings provide new insights into the regulation of secondary wall formation and its integration with environmental cues, advancing our understanding of grass stem development.

**SIGNIFICANCE:** Understanding how grasses build strong stems is essential for improving biomass production and crop resilience. In grasses, stem elongation and secondary cell wall thickening occur in distinct zones, yet the precise timing and regulation of this process remain unclear. To investigate this phenomenon, we developed a real-time imaging system to track the expression of *CESA8*, a key gene involved in cellulose synthesis. Our findings reveal that secondary wall thickening follows a daily rhythm controlled by temperature rather than light. These insights provide a foundation for optimizing plant architecture in bioenergy crops, improving their efficiency and sustainability.

## INTRODUCTION

In grasses, the growth of above-ground vegetative tissues occurs through repeated iterations of phytomers, each comprising a leaf blade, leaf sheath, stem internode and node, and an axillary bud. The intercalary meristems located at the node-internode junction at the base of each phytomer initiate and facilitate internode growth. As stem elongation begins, cells within these intercalary meristems undergo division, followed by elongation and maturation, each process occurring in distinct spatial regions characterized by specific molecular and cellular changes (1). While the precise location of the intercalary meristem is unknown, its progeny cells, which are actively dividing, remain relatively small and are surrounded by a primary cell wall, allowing for cell division (1). After elongation, some cell types develop a thick secondary cell wall that provides mechanical strength necessary for upright growth and water transport (2, 3). Grass leaves and stem internodes exhibit similar developmental gradients from base to tip, beginning with active cell division at the base followed by a zone of elongation and then maturation (4). As a result, growth at the base of each phytomer pushes older cells upward as they mature. The rate of this growth varies both throughout development and within a single day (5, 6).

In grass stem development, the transition from cell elongation to secondary cell wall thickening is characterized by a shift in gene expression as well as changes in cell wall composition (1). Primary walls are thin and flexible, and lack lignin, whereas secondary walls are substantially thicker, composed mostly of crystalline cellulose as well as hemicelluloses and lignin, making them rigid and less permeable (7). Secondary wall thickening in the stem is orchestrated by a network of mostly MYB and NAC family transcription factors which regulate the genes that encode enzymes involved in synthesis of cellulose, hemicelluloses, and lignin (7, 8). The large suite of genes associated with secondary wall synthesis including the cellulose biosynthesis associated genes *CELLULOSE SYNTHASE A4* (*CESA4*), *CESA7*, *CESA8*, *COBRA-like*, and *KORRIGAN*, the hemicellulose biosynthesis genes that can vary across eudicots and grasses, lignin biosynthesis genes like *4CL*, *CAD*, and *COMT*, have remarkably correlated gene expression pattern (9–12).

In the presence of photocycles and thermocycles, eudicot stem elongation oscillations can reach their peak at dawn (13, 14), dusk (15, 16), during daylight hours (17, 18), or at night (19), depending on the plant species, as well as tissue, and condition. These peak growth periods persist under constant temperatures when photocycles are present (14, 20) or under constant light conditions with thermocycles. Once established, these growth rhythms continue in the absence of external cues (21, 22). In contrast, elongation of grass stems and leaves is regulated solely and positively by temperature and not photocycles or the circadian clock (6, 14). Observations of transcript abundance in the grass *Brachypodium distachyon* revealed a similar pattern of influence on genes associated with the thickening of secondary cell walls (9). Genes associated with the biosynthesis of cellulose, hemicellulose, and lignin along with their associated transcription factors, exhibited a diurnal rhythm that was driven solely by temperature cycles.

Emerging leaves of *B. distachyon* elongate at a rate of 2 to 20 mm per day (6) while maize leaves at 2 to 3 mm per hour (14, 23). This is a phenomenon easily quantified throughout the day using various techniques (22, 24). In contrast, secondary cell wall growth occurs at a microscopic level, with mature interfascicular fiber cell walls measuring just a few microns in *B. distachyon* and rice stems (2, 25). Consequently, quantifying wall thickness of cells within a stem over the course of a single day is impractical. To investigate the cues regulating secondary cell wall thickening, we developed a luciferase reporter system that enables real-time measurement of the relative expression of *CESA8,* which was chosen because cellulose is the most abundant component of the secondary cell wall and of the *CESA* genes it is the most highly expressed. This reporter consists of the *CESA8 cis*-regulatory region fused to the fire-fly luciferase gene (*CESA8::LUC*). By capturing whole plant luminescent images, we were able to observe spatio-temporal expression patterns, contextualizing the results in relation to growth and development. In this study, we examined the effect of light, temperature, and the circadian clock on the *CESA8* expression as an indicator of secondary cell wall thickening.

## RESULTS

### *CESA8::LUC* is highly expressed during stem elongation in nodes and specific internode regions

To determine the location and timing of secondary wall thickening, we quantified bioluminescence in *B. distachyon* plants stably transformed with a *CESA8::LUC* reporter. In above ground tissues, we identified three areas with high luminescence: stem nodes, a specific region of elongating stem internodes, and the inflorescence (**Fig. 1A-C**). When separated from the stem, leaf and sheath displayed minimal expression (**Fig. S1**). Intriguingly, we found consistent patterns of localization within elongating internodes of the stem. Starting from the base of an internode, this pattern consisted of a low luminescence region referred to as the *CESA8* “dark zone”, approximately 3 mm in length above the node, and a *CESA8* “bright zone” with peak intensity approximately 5 mm above the node, which gradually diminished going up the internode. While the intensity of the node and bright zone changed throughout internode development **(Fig. 1D)**, the spatial patterning of the zones remained consistent with respect to distance from the node **(Fig. S2)**. These results suggest that immediately above the node, which is an intercalary meristem, cells are dividing and elongating, with minimal secondary wall thickening and where bioluminescence is high in the *CESA8* bright zone secondary wall thickening is high. Bioluminescence diminished around 12 mm above the node, presumably where secondary wall thickening is nearly complete and the stem reaches full maturity, at least with regard to cellulose accumulation within the secondary walls. While internodes can rapidly elongate, a node does not noticeably change in size, but *CESA8:LUC* expression remained relatively high in the early and middle stages of internode elongation **(Fig. 1E)**. This suggests that nodes are enriched for secondary walls. To test this, we conducted histological analysis and observed pronounced thickening of the vascular and interfascicular cells within the node (**Fig. S3**). Similarly, fully elongated spikelets exhibited very high *CESA8* expression. We performed a histology to investigate cell wall thickening of the seed which revealed areas of very thick walled cells in the lemma and palea surrounding the seed, but not in the seeds (**Fig. S4**). Thus, very thick secondary cell walls were observed in tissues where the *CESA8::LUC* reporter showed the highest luminescence.

**Fig. 1.**
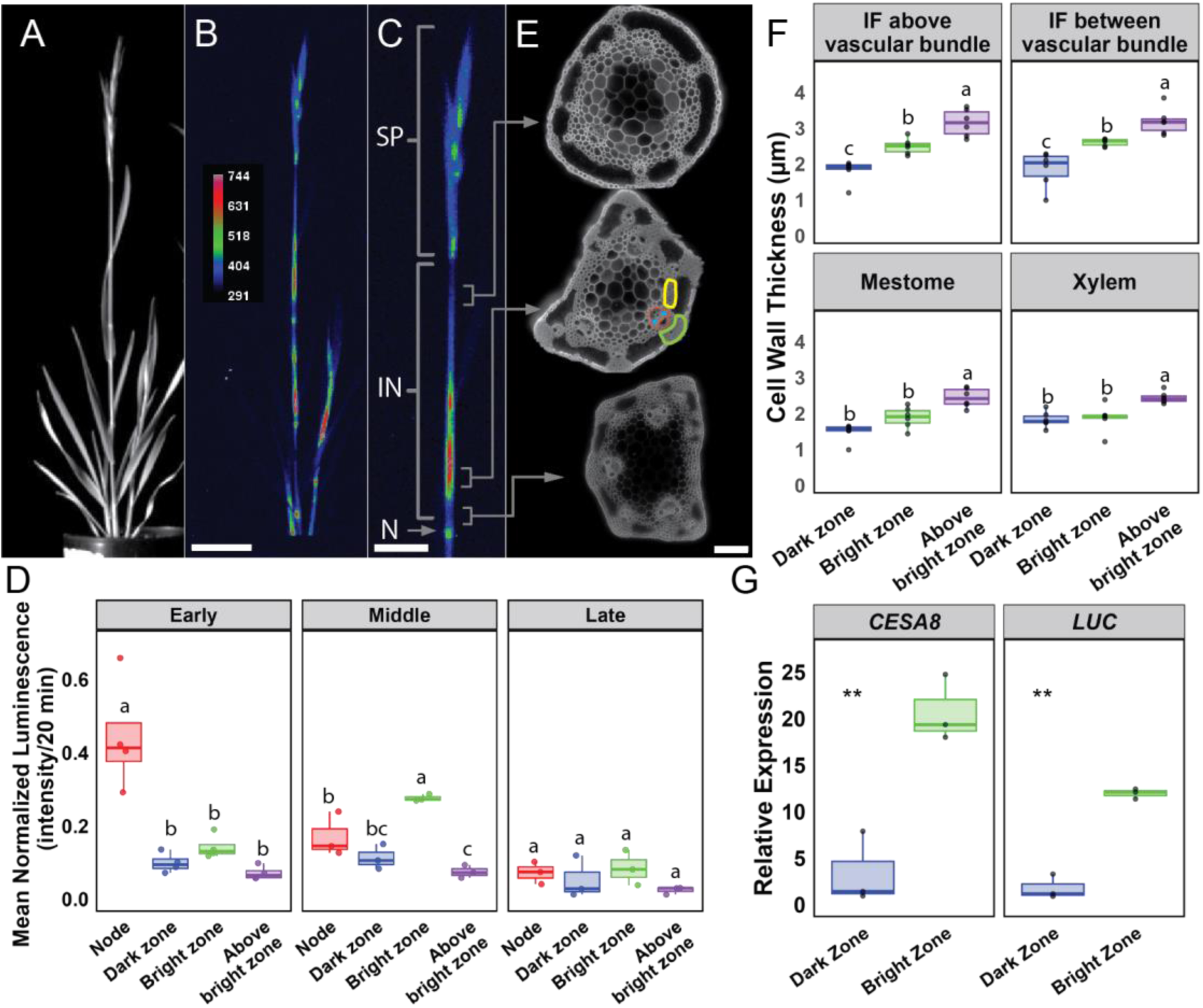
Spatial distribution and intensity of *CESA8::LUC* during stem elongation. **(A)** Bright field and **(B)** luminescence image of a *CESA8::LUC Brachypodium distachyon* plant captured with a CCD camera. The heat map represents luminescence intensity in arbitrary units, ranging from low (blue) to high (red). **(C)** In the expanded view of the top internode, luminescence is brightest in the inflorescence (SP), node (N), and near the bottom of the internode (IN). **(D)** Mean luminescence was quantified in each region, at three stages of development determined by top internode length (early: internode length < 2.5 cm, middle: internode length > 2.5 cm and < 4.5 cm, late: internode length > 4.5 cm). Regions are designated by distance from the peduncle bottom node: node < 1.5 mm, 1.5 mm < dark zone <5 mm, 5 mm < bright zone < 12 mm, above bright zone > 12 mm. N = 3 or 4 plants. **(E)** Fluorescence micrographs of Basic Fuchsin stained transverse internodes sections from above the bright zone (top), within the bright zone (middle), and in the dark zone (bottom). In the bright zone image, interfascicular cells located above the vascular bundles are circled in green, those between the vascular bundles are circled in yellow, mestome cells are highlighted in red, and xylem cells are highlighted in blue. Scale bars 100 µm. **(F)** Quantification of interfascicular fiber (IF), mestome, and xylem wall thickness. Groups that do not share a letter are significantly different using Tukey’s HSD test, *p* ≤ 0.05. **(G)** Relative expression of *CESA8* and *LUC* from three biological replicates using QRT-PCR and normalized relative to three control genes, Bradi1g25170, Bradi4g24887, and Bradi3g49600. Student’s t-test, \*\**p* ≤ 0.01.

### *CESA8::LUC* is coincident with *CESA8* mRNA and wall thickening

To investigate further whether *CESA8::LUC* bioluminescence correlates with the presence of secondary cell wall thickening, we performed a histological experiment in which we imaged *CESA8::LUC* plants and harvested tissue based on the luminescence localization. We used Basic Fuchsin staining of transverse stem internode sections to measure the secondary cell wall thickness of interfascicular fiber cells, mestome, and xylem cells (**Fig. 1E**). Interfascicular fiber cells are sclerenchyma cells found between vascular bundles, while mestome cells are a type of vascular bundle sheath cell. The walls were thinnest at the base of the stem internode, the dark zone, increasing in thickness and stain intensity in the bright zone (**Fig. 1F**). In the bright zone, more secondary wall deposition was observed with the exception of vascular bundle related cell types. Xylem and mestome cell wall thickness did not increase from dark zone to bright zone, and only slightly increased in above bright zone tissue indicating that they mature earlier than interfascicular fiber cells.

To characterize the secondary cell wall cellulose, tissue from the dark zone, bright zone, and area above the bright zone was harvested, dried, and ground into a fine powder for X-ray diffraction. We estimated cellulose crystallinity using a peak deconvolution method, analyzing X-ray diffraction intensities from five crystalline peaks, including the characteristic peak at 22.6° **(Fig. S5)**. Crystallinity increased most from the dark zone to the bright zone, with a slight additional increase observed from the bright zone to the region above it. Additionally, mRNA was extracted from these tissue regions for RT-qPCR analysis, revealing that our reporter gene, *CESA8::LUC*, and *CESA8* mRNA expression levels directly correlate with each other and with the luminescence data **(Fig. 1G)**. These results indicate that luminescence from the *CESA8::LUC* transgene effectively reflects native *CESA8* transcript levels and serves as a reliable marker of secondary wall thickening.

### Thermocycles alone affect daily rhythm of *CESA8* expression

Contrary to eudicots, in grasses such as maize, rice, and *B. distachyon* diurnal shoot growth is driven by thermocycles, but not photocycles (5, 6, 14). To investigate whether daily rhythms of secondary cell wall thickening are similarly regulated by thermocycles alone, we performed a series of time lapse imaging experiments with or without photo and thermocycles using our *CESA8::LUC* reporter line. *CESA8::LUC* plants were imaged following when the awns first emerge from the flag leaf, awn tipping, as the peduncle elongates. All plants were entrained in the imaging chamber in a long day with photo and thermocycles (light dark hot cold, LDHC) for two days and then released and monitored every 3 hours for 4 days in experimental conditions. Mean pixel intensity for each plant during each time point was measured (**Fig. 2 and Fig. S6**).

**Fig. 2.**
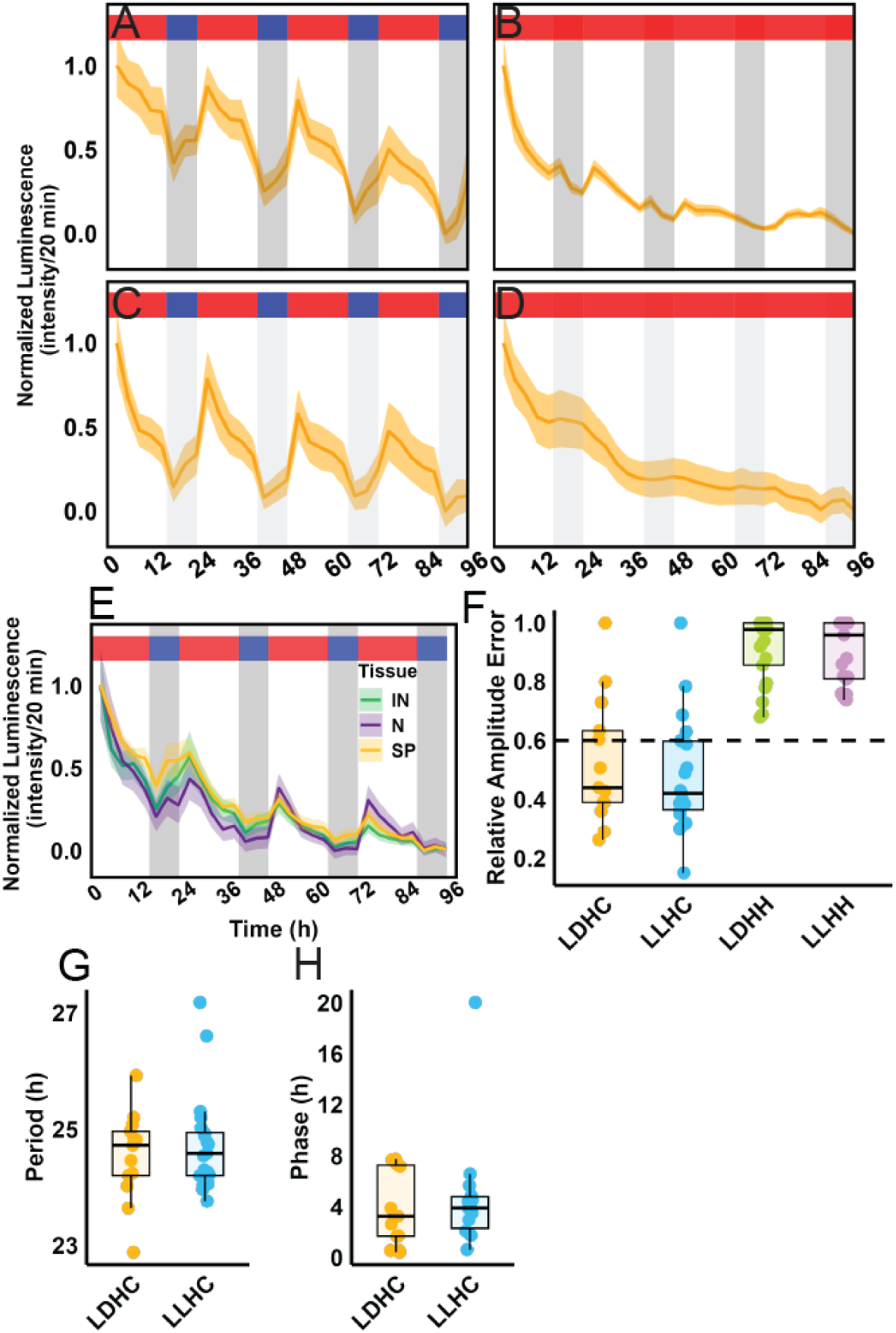
Diurnal thermocycles regulate the morning-phase expression of *CELLULOSE SYNTHASE A8*. *CESA8::LUC* plants were entrained under photo and thermocycles and then imaged under **(A,E)** photo and thermocycles, **(B)** constant light with thermocycles, **(C)** constant 25°C with photocycles or **(D)** constant light and 25°C. **(E)** Luminescence of inflorescence (SP), node (N), and near the bottom of the internode (IN). Circadian **(F)** rhythmic amplitude (measured as RAE) **(G)** period length, and **(H)** phase were determined using BioDare. Luminescence data are means ± SEM, n= 7. Dark gray shading represents night, light gray shading represents subjective night, red shading indicates 25°C, and blue shading indicates 17°C. The experiments were performed at least twice with similar results (Fig. S7). LDHC, photo and thermocycles; LDHH, photocycles and constant temperature; LLHC, thermocycles and constant light; LLHH, constant light and constant warm temperature.

Similar to previous studies of grass elongation growth, we observed that rhythmic expression of *CESA8::LUC* is driven by thermocycles and not photocycles or the circadian clock. A majority of the plants had a relative amplitude error (RAE) less than 0.6, indicating a significant daily rhythm (**Fig. 2F**). In both time course experiments with a thermocycle, *CESA8* exhibited a period of approximately 25 h (**Fig. 2G**). On the other hand, all of the plants monitored without a thermocycle (LDHH and LLHH) had an RAE greater than 0.6, indicating a lack of rhythmicity. *CESA8* exhibited an early morning phase, peaking approximately 3 h into the subjective day, followed by a decrease in expression during 25°C days and an increase during 17°C nights (**Fig. 2H**). These results suggest that there is no circadian clock driven anticipation of daily changes in light or temperature. Furthermore, prolonged daytime temperatures repress *CESA8* expression and subsequently secondary wall thickening, while prolonged cooler nighttime temperatures activate *CESA8*.

### Diurnal pattern of *CESA8* expression is consistent across tissue type

In previous analyses we measured whole plant luminescence to determine if *CESA8::LUC* expression was rhythmic in response to specific diurnal cues. This approach however, overlooks the possibility that different tissues have different expression patterns. To investigate this possibility we analyzed the expression pattern of specific tissue types: nodes, internodes, and spikelets. The waveform was similar across tissue types with a daily rhythm peaking in the morning (**Fig. 2E**). This consistent daily rhythm across different areas of the plant suggests that the observed patterns in *CESA8::LUC* expression throughout the day are synchronized by thermocycles.

### Initial change in *CESA8* expression is positively correlated with temperature

To investigate whether warm temperatures repress *CESA8* expression or cold temperatures activate it, we conducted a series of temperature experiments involving either a temperature pulse or a transition to shorter or longer day thermocycles. Under standard thermocycles, a 3 h daytime cold pulse of 17°C caused a sharp decrease in *CESA8* expression, which then returned to levels comparable to an uninterrupted thermocycle (**Fig. 3A and Fig. S7**). However, the typical nighttime increase in *CESA8* expression during prolonged cold exposure remained unaffected by the daytime cold pulse. Conversely, a 3-hour nighttime warm pulse at 25°C elevated *CESA8* expression to typical morning levels, followed by a decline when the temperature reverted to 17°C (**Fig. 3B and Fig. S7**). These findings indicate that *CESA8* expression responds rapidly to temperature fluctuations, with warm temperatures promoting expression and cold temperatures suppressing it.

**Fig. 3.**
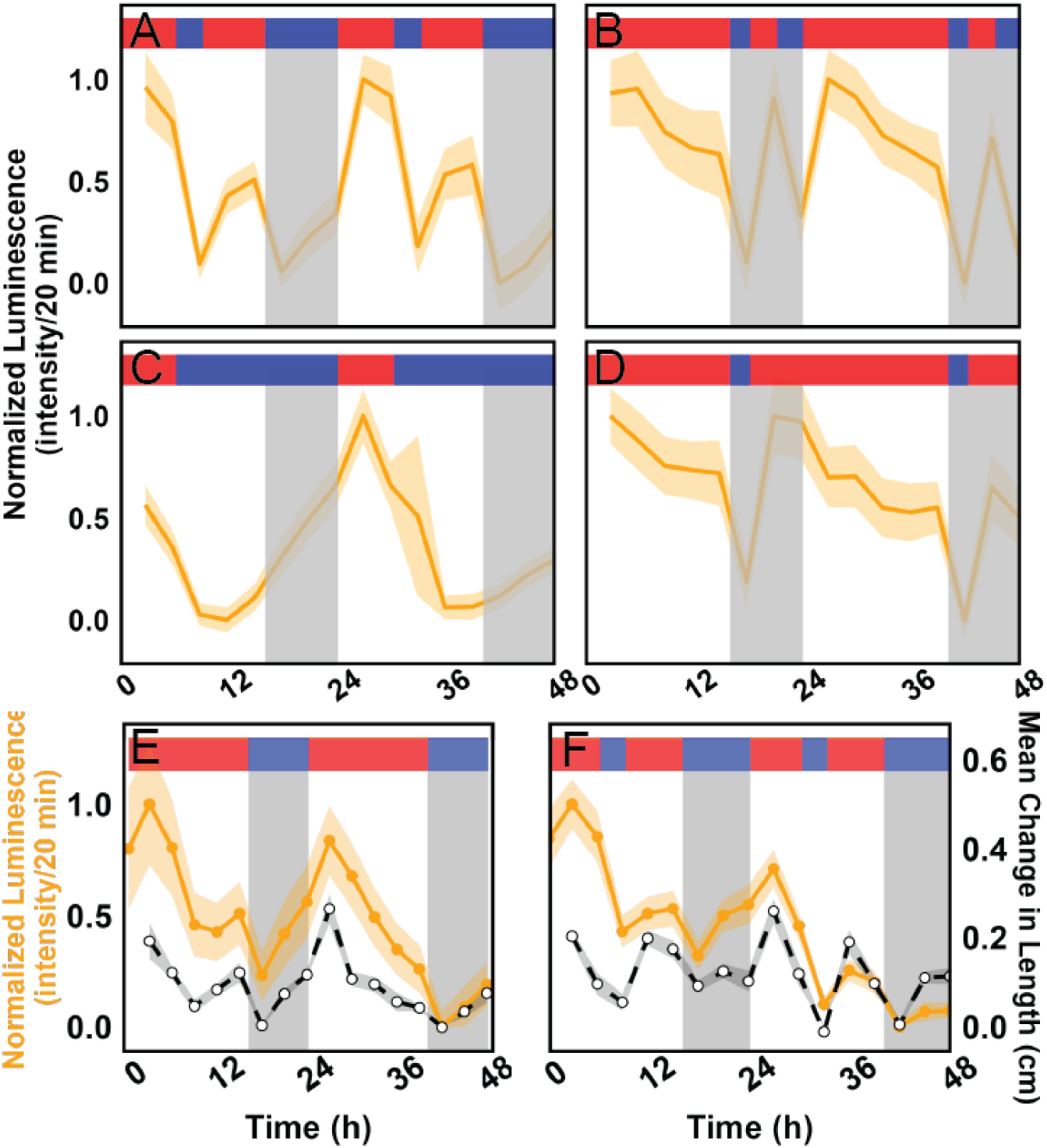
*CELLULOSE SYNTHASE A8* expression responds immediately to temperature changes, increasing with warmth and decreasing with cold. *CESA8::LUC* plants were entrained under photo and thermocycles, transferred to thermocycles alone and then given an **(A)** 17°C cold pulse for 3 h midday or a **(B)** 25°C warm pulse for 3 h at night. Long day entrained plants encountered **(C)** early nighttime temperature, equivalent of a short day thermocycle (6 h 25°C:18 h 17°C) or **(D)** early daytime temperature, equivalent to very long day thermocycle (21 h 25°C:3 h 17°C). **(E)** Luminescence and elongation of the primary stem were measured for each timepoint of plants grown with a photo and thermocycle like Fig. 2A or **(F)** daytime cold pulse like (A). Luminescence data are means ± SEM, n=7. The experiments were performed at least twice with similar results.

To examine the prolonged effect of temperature on *CESA8* expression, we performed a series of extended pulse experiments where the previous pulse was extended into the next temperature cycle, resulting in either early-night or early-day. Under early-night thermocycles, plants exhibited a pattern similar to long day conditions, with *CESA8* expression gradually rising during the cooler night and decreasing during the warmer days (**Fig. 3C and Fig. S7**). Under early-day conditions, *CESA8* expression quickly returned to morning levels before resuming typical diunral rhythm of decreasing throughout the day (**Fig. 3D and Fig S7**). These findings suggest that while *CESA8* follows a diurnal thermocycle, rising at night and falling during the day, its immediate response to a temperature shift is distinct and follows an inverse pattern.

### Stem elongation is directly correlated with *CESA8* expression

With both elongation and *CESA8* expression being driven by temperature we sought to investigate whether they are directly correlated. In luminescence data from both LLHC and LLHC with a temperature pulse we measured the elongation of each plant’s main stem as well as that main stem’s luminescence (**Fig. 3E-F**). Results from this analysis show that elongation and *CESA8* expression are correlated with a Pearson correlation coefficient of 0.82 and 0.58 for LLHC and LLHC cold pulse, respectively.

### An incoherent feed-forward loop model captures *CESA8* dynamics

To explain our results, we developed a simple mathematical model centered on *CESA8* dynamics. The model accounts for the synthesis and degradation of *CESA8* mRNA (variable *X*) and posits that temperature (T) has a dual effect on its expression rate: a direct activation and an indirect inhibition mediated by a temperature-sensitive factor (variable *S*) (**Fig. 4A-B**). This incoherent feed-forward loop generates a transient pulse of *CESA8* expression at the onset of the warm period, before *S* accumulates sufficiently to repress it (26, 27). The model qualitatively reproduces the experimentally observed dynamics under thermocycles, including the 3 h phase shift relative to the onset of the warm period and the apparent inverse relationship when a warm pulse is applied during the cold night or a cold pulse during the warm day (**Fig. 4C-H**). To assess the robustness of the model across parameter variations, we conducted simulations in which the value of each kinetic parameter was randomly increased or decreased by up to 10%. Twenty simulations were performed, each with a new generated parameter set. These results indicate that while variability primarily affects the amplitude and, to a lesser extent, the phase of the response, the overall dynamics remain consistent across simulations.

**Fig. 4.**
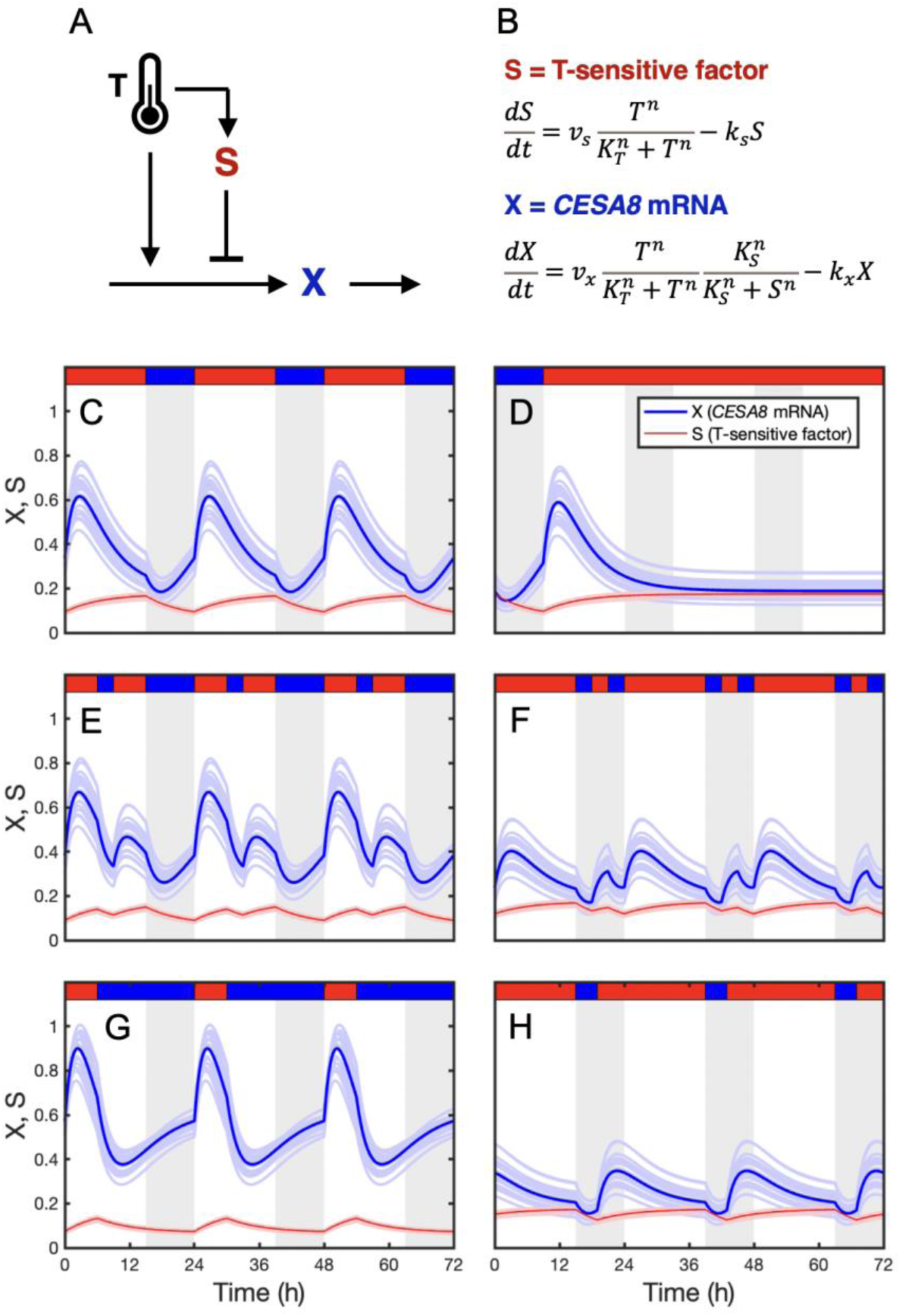
The dynamics of *CESA8* can be reproduced by an incoherent feed-forward loop model. **(A)** In this model, temperature (T) directly induces the expression *CESA8* (variable *X*) while also indirectly suppressing it through a hypothetical temperature-sensitive factor (variable *S*). **(B)** The activation of S and X expression by temperature (T), along with the inhibition of X expression by S, follows Hill function dynamics. Numerical simulations of *S* (red) and *X* (blue) over time under **(C)** thermocycles, **(D)** release to continuous daytime conditions, **(E)** daytime cold pulse, **(F)** nighttime warm pulse, **(G)** early nighttime temperature, **(H)** early daytime temperature show qualitative agreement with the experimental observations. Parameter values: *v_X_* = 1.2 h^-1^, *K_T_* = 23°C, *K_S_* = 0.1, *n* = 4, *k_X_* = 0.36 h^-1^, *v_S_* = 0.045 h^-1^, *k_S_* = 0.15 h^-1^, *T_warm_* = 25°C, *T_cold_* =17°C. The light blue and red curves represent results obtained by introducing variability in parameter values. Specifically, the values of *v_x_*, *K_T_*, *K_S_*, *k_x_*, *v_s_*, and *k_s_* were randomly increased or decreased by up to 10% from their default values. A total of 20 simulations were performed, each using a new randomly generated set of randomized parameter values.

## DISCUSSION

Plant sources dominate global biomass, contributing approximately 80% of the total 550 gigatons on Earth (28). Other estimates suggest that the lignocellulosic proportion of grasses, i.e., secondary cell walls, constitutes 60%-80% of dry plant matter (29). Given that plant cell walls are a larger portion of planetary biomass than animals, bacteria, and archaea combined, it is surprising how little is known about the timing of secondary wall formation. Indeed, tracking cell wall thickening is far more challenging than measuring elongation and expansion, as root and leaf length can be easily observed relative to secondary wall thickening which is just a few micrometers wide and is obscured from observation inside the plant. Thus, to investigate this, we developed a bioluminescent reporter system for real-time, non-destructive measurement of whole-plant *CESA8* transcription.

Transcript levels of secondary wall-associated CESAs are elevated in tissues undergoing secondary wall thickening, particularly in the developing stem (12, 30). Many key transcriptional regulators of cellulose biosynthesis were initially identified through transcription profiling (9–12, 31, 32), highlighting the importance of gene expression timing and abundance in secondary wall formation. The *CESA8::LUC* transgene was designed to serve as a reporter of native *CESA8* transcript and as an indicator of secondary wall thickening. When fused to the *CESA8* coding sequence, the *CESA8 cis*-regulatory region can genetically rescue the stunted culm growth and reduced cellulose phenotype of the *cesa8-1* mutant (33). To validate *CESA8::LUC* bioluminescence as a reporter of *CESA8* expression, we measured *LUC* and *CESA8* transcript levels across different regions of the developing internode. The results revealed a strong correlation between *LUC* and *CESA8* expression dynamics, consistent with *B. distachyon* time-course data and gene atlas findings (9, 12, 30, 33). Additionally, we found that secondary wall thickness closely aligns with *CESA8::LUC* reporter activity. At the base of the internode prior to *CESA8::LUC* detection, where cells originate from the intercalary meristem, interfascicular fiber walls were thinnest. In the bright zone, where *CESA8* expression peaks, wall thickness increased and continued beyond this region. As expected with increased secondary wall deposition, cellulose crystallinity rose sharply above the intercalary meristem. Further analysis of two other tissues with bright bioluminescence, stem nodes and spikes, revealed that both also possessed extraordinarily thick secondary walls. Together, these findings demonstrate that our reporter reflects *CESA8* transcription and serves as a reliable indicator of secondary wall thickening.

While our findings, along with those of others, support a relationship between *CESA8* transcript abundance and secondary wall cellulose synthesis, it is critical to consider the role of post-translational regulation in modulating cellulose synthase complex (CSC) activity. The rosette-shaped complex responsible for primary wall synthesis has been observed to persist within the Golgi complex for more than 48 hours following the inhibition of protein synthesis by cycloheximide treatment. Notably, its longevity is inversely correlated with temperature increases (34). This extended Golgi residency suggests that circadian fluctuations in gene expression are unlikely to directly dictate cellulose synthesis rates over short timescales. However, it is essential to recognize that the CSC is only catalytically active at the plasma membrane, not within the Golgi (35). In *A. thaliana*, primary wall CSCs remain active in the plasma membrane for a remarkably short duration, approximately 7 to 8 minutes, before undergoing clathrin-mediated endocytosis (36–38). In cotton fibers, the CESA8 ortholog, GhCESA1, exhibits a half-life of less than 30 minutes and is undetectable after 4 hours (39). This broad yet seemingly disparate body of observations underscores the inherent challenges associated with directly assessing CESA protein activity, particularly in developing green tissues such as grass stems. The interplay between transcriptional and posttranscriptional regulation, CSC trafficking, and post-translational modifications remains a critical frontier in understanding spatiotemporal control of cellulose biosynthesis in diverse plant systems (40–44).

The *CESA8* reporter signal was greatest in the stem nodes and internodes and the spikelets. Unlike the node and spikelet, the internode is highly dynamic, with the peduncle growing to approximately 7-8 cm in our experiments. The *CESA8* internode bright zone occured at a fixed distance from each node, beginning approximately 5 mm above the base of the internode. Notably, this region is enclosed within the sheath, which likely provides additional mechanical support. Transcription profiling of the bottom 1 cm of an elongating sorghum internode was not enriched for genes involved in secondary cell wall synthesis (1) which aligns with our observations of minimal *CESA8* expression in a comparable region of *B. distachyon*. However, tissue immediately above the base of the sorghum internode was not assayed, precluding a direct comparison to localization of *CESA8* expression in our study. These findings highlight a spatial coordination of cell wall synthesis as a critical component of grass stem development, maintaining structural integrity during rapid growth and elongation.

The timing of rapid elongation in eudicots is regulated by diurnal changes in light and temperature, and the circadian clock, whereas in grasses, the daily rhythm of leaf and stem elongation is primarily regulated by temperature alone (6, 14, 20, 45, 46). Our findings, along with previous research, demonstrates that in *B. distachyon*, the transcription of secondary cell wall thickening genes follows a diurnal rhythm driven by thermocycles (9). Under a diurnal thermocycle, warm daytime temperatures resulted in a decrease in *CESA8* expression, followed by a steady increase during the cooler nighttime period. However, no detectable rhythmicity was observed under free run conditions. Different organs of *A. thaliana* exhibit variability in circadian clock speed, but this variation is significantly reduced under a photoperiod (47). Our analysis of internodes, nodes, and spikelets revealed no differences in period or phase under photocycles and thermocycles together. Unlike the thermocycle-driven rhythm of *CESA8*, which was inversely correlated with temperature change, the reporter exhibited the opposite response to a temperature pulse. Similar to elongation rate, *CESA8* expression was positively and highly responsive to temperature. Previous studies have shown that the stem elongation rate of *Lilium longiflorum* decreases from start to end of a light period and increases from start to end of a dark period and that a cold pulse at the start of a light period reduces elongation (48, 49). Within a three-hour window, we observe that both elongation and *CESA8* expression decreased in response to cold pulses or increase in response to warm pulses. However, extended pulses revealed an opposite response in *CESA8* expression, suggesting the interplay of both rapid temperature responsiveness and temperature-driven clock regulation.

The time profiles observed under different thermocycles can be qualitatively reproduced by a simple mathematical model incorporating a feed-forward loop motif. In this model, temperature directly activates and indirectly inhibits *CESA8* expression. Consistent with experimental observations, the model predicts a delayed inversion in *CESA8* at the cold-to-warm transition, accompanied by a transient surge. This transient peak may serve as a signal marking the start of the day and potentially extending growth during this early phase.

By leveraging a real-time bioluminescent reporter system, we have demonstrated that *CESA8* transcription follows a diurnal rhythm driven by thermocycles, with expression increasing during cooler nighttime periods and decreasing during warmer daytime conditions. Our results further suggest an interplay between rapid temperature responsiveness and a thermocycle driven clock with regards to *CESA8* expression and elongation. This study provides novel insights into the spatial and temporal coordination of secondary cell wall synthesis in grasses, highlighting its role in maintaining structural integrity during rapid stem elongation. Future research exploring the molecular mechanisms linking temperature signaling to transcriptional regulation will further enhance our understanding of grass development and biomass accumulation.

Grass stem development differs from that of eudicots, relying on stacked intercalary meristems located at the base of internodes rather than extension of apical and lateral meristems. As a result, multiple positions within the growing stem undergo active cell division and elongation before structural reinforcing through secondary cell wall thickening. A drawback of this growth strategy is that intercalary meristems must withstand substantially greater mechanical stress than cells derived from an apical meristem. Our findings highlight the complex relationship between temperature-regulated stem elongation and secondary cell wall synthesis in grasses. Internode elongation increases with warming temperatures, creating a greater demand for wall thickening. Consistent with this model, a decrease in temperature slows both elongation and *CESA8* expression. The apparent paradox of diurnal temperature cycles having the opposite effects may be explained by the role of daily rhythms in preparing the plant for the anticipated demands of the coming warmer day.

## MATERIALS AND METHODS

### Development of Bioluminescent Imaging Chamber

The bioluminescence imaging chamber consists of an inner and outer shell. The inner shell of a 61 x 122 x 183 cm grow tent (Gorilla Grow Tent, Santa Rosa, CA) placed on its side on a 183 x 61 x 214 cm metal wire shelving unit (H-2948-86, ULINE) with two wire shelves at a height of 80 and 205 cm. An outer shell of Black Out Heavy 6 Mil Plastic (B & H Foto & Electronics Corp., New York NY) covers the entire shelving unit. We removed a 45 x 19 cm section from the top shelf to mount the custom light-emitting diodes (LED). The light array consists of royal blue (448 nm), green (530 nm), red (627 nm), deep red (655 nm), far red (720 nm), and 6500 K and 5650 K cool white (**Table S1**). Diodes were mounted onto a grade 6063 aluminum heat sink from a Makers LED Grow Light Kit (LEDSupply, Randolph, VT). The lights are mounted into the lining of the inner and outer shell. This design allowed for the heat sink to be exposed outside the chamber to reduce heat transfer from the LEDs into the chamber. Temperature control is accomplished with a GE 6100-BTU Portable Air Conditioner (Part number 2759118, Lowe’s), wiring in a new thermostat (Stego Smart Sensor temperature/humidity sensor, part number 014202-00, AutomationDirect.com) fixed to the inner shell adjacent to where plants are staged. Control of the light and temperature cycles is achieved through a CLICK Programmable Logic Controller (ClickPLUS CPU, C2-01CPU). Temperature and light cycles are programmed on CLICK Programming Software version 3.40. An Andor iKon-M 934 CCD camera (Oxford Instruments, Concord MA) with a Nikkor 50 mm f /1.2 lens was used to capture plant bioluminescence as described below. NIS-Elements D 5.30.04 was used to control capture settings and manage time-lapse imaging.

### Plant Materials and Growth Conditions

The *CESA8::LUC* reporter plasmid was developed by recombining the codon optimized *LUC2* luciferase (*LUC*) coding sequence into the *B. distachyon CESA8* Utility Promoter Vector (Genbank KT962835) (33, 50). The *CESA8* promoter fragment includes the 1,888 nucleotides of sequence upstream from the translational start site of the *CESA8* gene of accession Bd21-3, BdiBd21-3.2G0638500. While not used here, the Bd21 reference genome gene identifier is Bradi2g49912 (30). The complete sequence and map of the plasmid are described in **Fig. S8**. The *CESA8::LUC* plasmid was transformed into *Agrobacterium tumefaciens* strain AGL1, which was then used for transformation of Bd21-3 embryonic callus as previously described (30). Plants harboring the *CESA8::LUC* reporter were regenerated from callus tissue and putative transgenic plants were initially confirmed by selection on 40 mg/L hygromycin. Genomic DNA was extracted from leaf tissue of putative transgenic plants and the *LUC* coding sequence was PCR-amplified using primers *Luc_F* and *Luc_R* (**Table S2**). Plants were also confirmed as transgenic by PCR and screening for bioluminescence.

Wildtype Bd21-3 and *CESA8::LUC* transgenic seeds were imbibed for 21 days at 4°C and then planted in Pro-Mix BX Mycorrhizae potting mix. The seeds were planted 1.5 cm deep in 22 cm tall Cone-tainer pots with a 3.8 cm diameter (Ray Leach SC10R, Stuewe & Sons, Inc.). Plants were first grown in a PGC-105 growth chamber (Percival Scientific, Perry, IA) with a ∼450 μmol·m^−2^·s^−1^ day period of 16 h at 25°C and a dark night period of 8 h at 17°C and watered daily. At the beginning of peduncle elongation, approximately BBCH stage 5.3 (51), plants were moved to the imaging chamber at the same long day thermo and photocycles with a daytime light intensity of ∼150 μmol·m^−2^·s^−1^. Consistent application of luciferin was achieved with the use of an automated, fine droplet mister (v5.0 Starter Misting System, MistKing, Ontario, Canada) that sprayed the plants with a 1 mM D-luciferin (Gold Biotechnology, St. Louis, MO) solution for 3 s, every 3 h.

Plants were allowed to acclimate to the imaging chamber and dissipate the effects of luciferase build-up by spraying with luciferin and delaying image acquisition 1 d. Images were then captured every 3 h for 4 d. To avoid the effects of delayed fluorescence, image capture began 5 min after the lights were turned off. Raw imaging data was processed using Fiji (52) to generate masks for each image, overlay the mask on the images, and apply a minimum threshold of 350 AU to eliminate background noise. To measure luminescence, regions of interest were manually drawn using Fiji. The raw luminescence values were then subtracted from the moving mean and divided by the moving standard deviation to normalize between 0 and 1. To measure elongation, a segmented line with a width of 3 pixels was drawn on the main stem of each plant in each image. Elongation was calculated by subtracting the length at the previous time point, three hours earlier. The mean elongation and standard error were then calculated for each time point.

### Histology of Stem Cell Walls

The peduncle of the tallest stem was excised when they were between 3 and 3.5 cm long. Nodes and seeds were collected for sectioning after the plants had fully senesced. The internode was segmented into three sections: the bottom 5 mm (dark zone), a 5 mm region above the dark zone (bright zone), and the uppermost 5 mm. The segments were embedded, sectioned transversely at 60 μm, and the wall thickness and staining intensity of either phloroglucinol-HCl or Basic Fuchsin were quantified as previously described (McCahill et al. 2024). The node and seed sections were embedded, sectioned transversely at 45 μm and 60 μm, respectively. Brightfield imaging of phloroglucinol-HCl sections was conducted using an Eclipse E200MV microscope (Nikon) equipped with a Pixelink 3 MP camera. For Basic Fuchsin stained sections, both fluorescence and brightfield imaging were captured using a BZ-X800 fluorescent microscope (Keyence). Fluorescent images were captured with an excitation wavelength of 545 nm and an emission wavelength of 605 nm. Five cross-sections from each of the three regions of five plants were analyzed using the line tool in Fiji and the average of cell wall thickness was calculated for each cross-section measured. In each cross section the walls of 7 mestome, 4 metaxylem, and 5 interfascicular fiber cells were quantified.

### Statistical analysis

Histological data were analyzed using analysis of variance and pairwise differences were assessed using Tukey’s HSD test with a significance threshold of *p* < 0.05. Biodare2 was used with the MESA algorithm, applying amplitude and baseline detrending, to calculate period and phase values for the time courses (51). Period and phase were reported only for plants considered rhythmic with an RAE below 0.6 (53).

### X-Ray Diffraction Profiling

X-ray diffraction and calculation of a crystallinity index were done as described by (30) with slight modifications. Peduncle stem tissue from the dark zone, bright zone, and above bright zone were harvested at stage 5.3 on the *B. distachyon* BBCH-scale. For each pool of tissue > 100 plants were harvested to attain a sufficient quantity for analysis. Stems were then dried for 2 weeks at 60°C in a HERA THERM Oven (Thermo Scientific). Tissue was ground using 3.2 mm diameter stainless steel metal balls (Biospec Products, Bartlesville, OK) in a ball mill (Mixer Mill, MM400, Retsch, Newtown, PA). Diffraction was analyzed on an X’Pert Pro powder X-ray diffractometer (PANanalytical BV, The Netherlands) operated at 45 kV and 45 mA using CuKα radiation at both Kα1 (λ = 1.5406 Å) and Kα2 (λ = 1.5444 Å). The diffraction profile was acquired from 5 to 40° in 0.0167 steps, with 66 s per step. The crystallinity index was calculated using the peak deconvolution method in MatLab, based on five Gaussian crystalline peaks. It was determined as the ratio of the combined area of all crystalline peaks to the total area under the curve (54).

### Measurement of transcript abundance

Total RNA was extracted using a kit (Plant RNeasy, Qiagen, Valencia, CA, USA) according to the manufacturer’s instructions. The peduncle internode of the main stem was harvested at stage 5.3 on the *B. distachyon* BBCH-scale (51). The peduncle was then subdivided into dark zone (1.5 to 5 mm above the node), bright zone (5 to 12 mm above the node), and then frozen in liquid nitrogen. Five individual plants were pooled together for each replicate. First strand cDNA was synthesized from 1 μg of total RNA using QuantiTect Reverse Transcription Kit (Qiagen). Three technical replicates of three biological replicates of QRT-PCR was then carried out using a QuantStudio 3 Real-Time PCR System. The expression values were normalized according to three control genes *Bradi1g25170*, *Bradi4g24887*, and *Bradi3g49600* identified as having low variance in transcript abundance across the developing stem. All oligonucleotides are described in **Table S1**.

## ACKNOWLEDGEMENTS

The microscopy data was gathered in the Light Microscopy Facility and Nikon Center of Excellence, and equipment constructed in the Advanced Digital Design and Fabrication Core Facility at the Institute for Applied Life Sciences, UMass Amherst with support from the Massachusetts Life Sciences Center. We thank Eva Farre for critical reading of the manuscript and helpful suggestions.

---

**Fig. S1.**
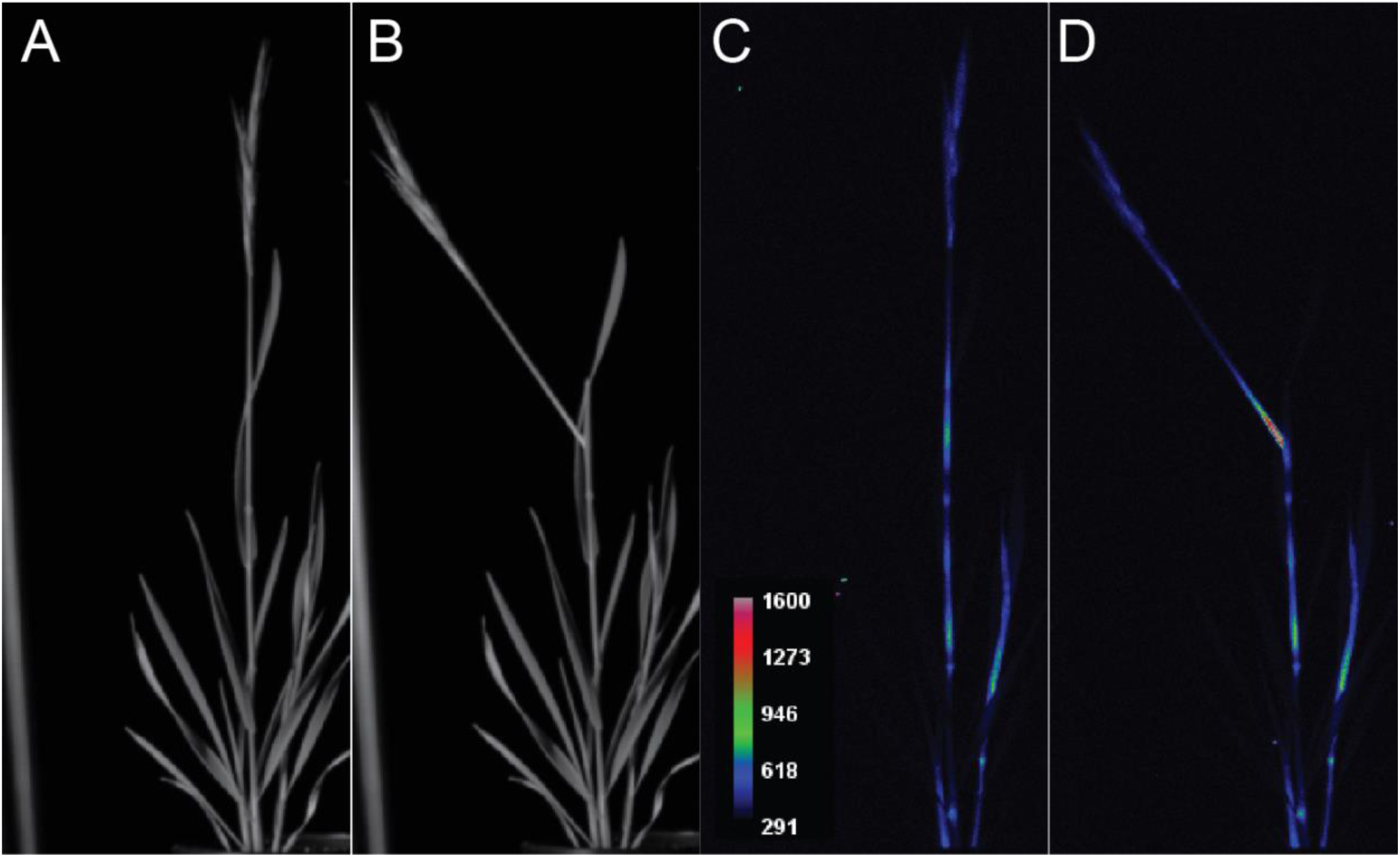
The leaf sheath does not contribute appreciably to luminescence. Spatial distribution and intensity of *CESA8::LUC* during stem elongation. **(A, B)** Bright-field images and **(C, D)** corresponding luminescence images captured with a CCD camera. The heat map represents luminescence intensity in arbitrary units, ranging from low (blue) to high (red). **(B, D)** Separation of the sheath from the stem internode reveals minimal detectable luminescence in the sheath.

**Fig. S2.**
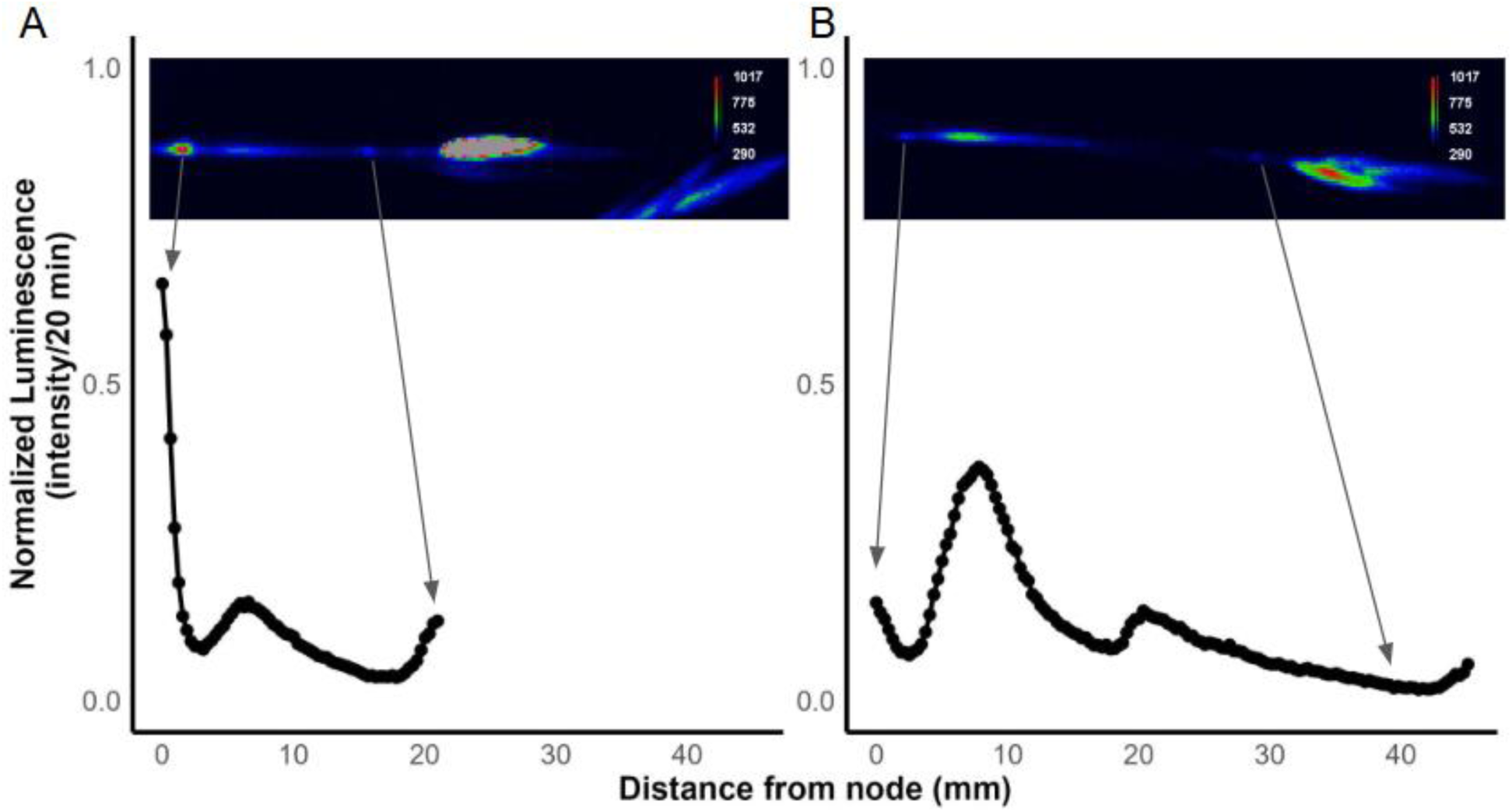
The peak position of the CESA8 luminescence zone remains constant relative to the node below. Normalized luminescence intensity across a single peduncle. **(A)** Two days after the internode elongation began and **(B)** three days later. The heat map represents luminescence intensity in arbitrary units, ranging from low (blue) to high (red).

**Fig. S3.**
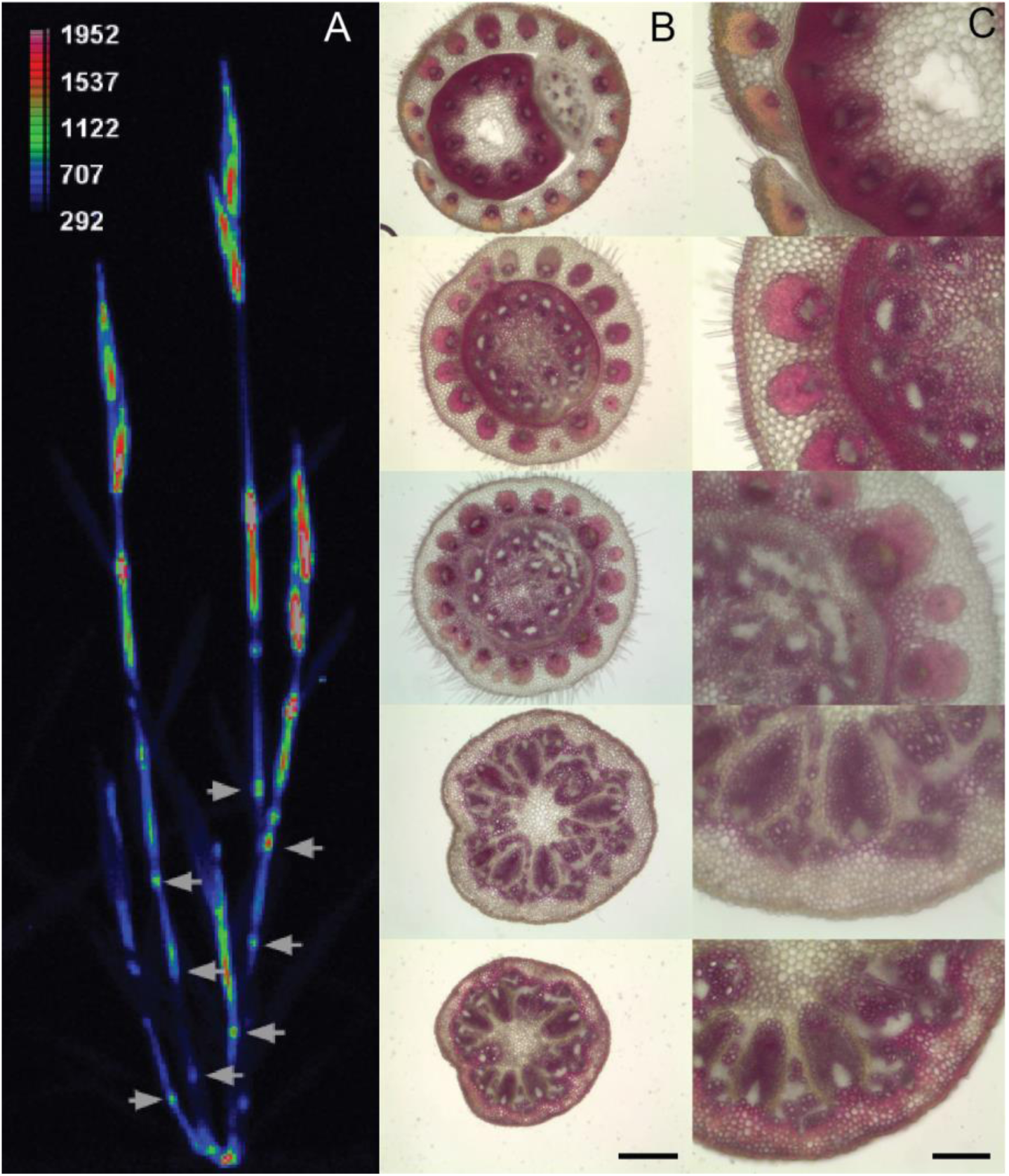
Histological characterization of the *Brachypodium distachyon* bottom peduncle node. **(A)** Representative image of above ground *CESA8::LUC* luminescence with arrows indicating the stem nodes. Warmer colors indicate greater luminescence. The heat map represents luminescence intensity in arbitrary units, ranging from low (blue) to high (red). **(B)** Phloroglucinol-HCl stained transverse sections of a fully senesced stem node, progressing upward from the base of the node to the point where the sheath separates from the internode, shown at 4x **(B)** and 10x **(C)** magnification. Scale bars 75 µm (A) and 30 µm (B).

**Fig. S4.**
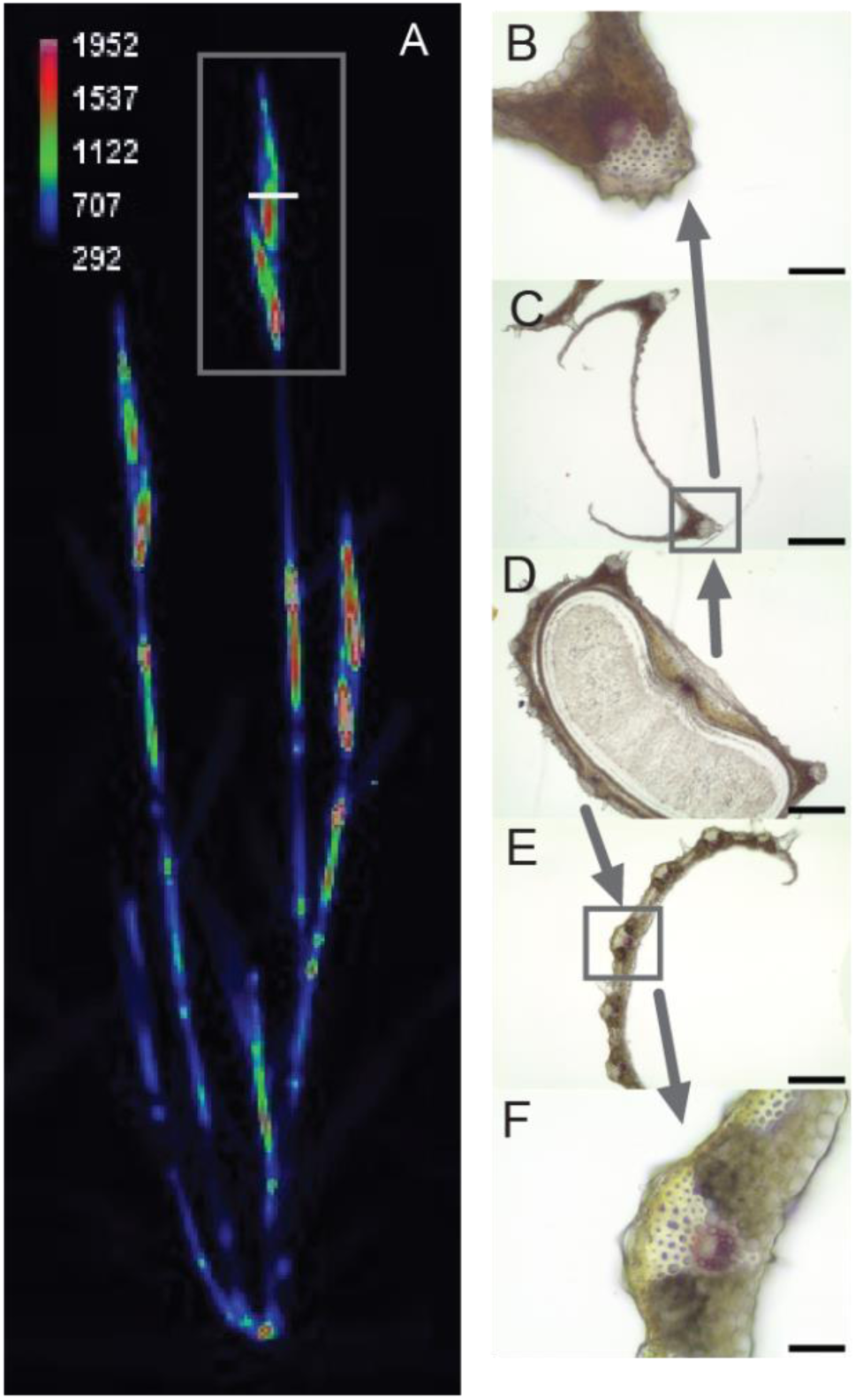
Histological analysis of *Brachypodium distachyon* seeds reveals extensive secondary cell wall thickening in the lemma and palea. **(A)** Representative image showing above-ground *CESA8::LUC* luminescence; the white line indicates the position of transverse sections. The heat map represents luminescence intensity in arbitrary units, ranging from low (blue) to high (red). Seeds with the palea **(B, C)** and lemma **(E, F)** intact were sectioned and stained with phloroglucinol-HCl, shown at 20X **(B, F)** and 4X **(C–E)** magnification. **(D)** Whole seed with both lemma and palea intact. Scale bars: 50 µm for 20X images, 250 µm for 4X images.

**Fig. S5.**
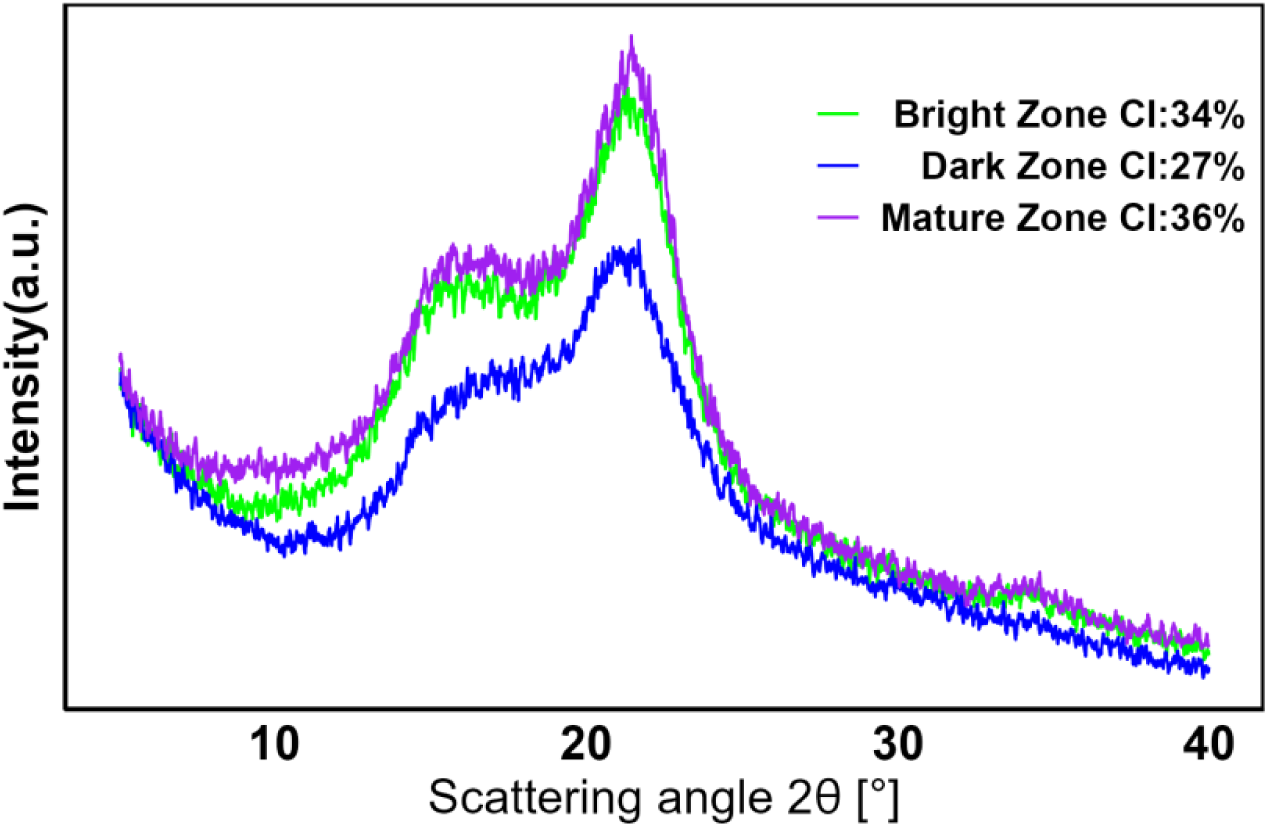
The crystallinity index of the stem internode increases from the base to the top. Spectroscopic analysis of cellulose crystallinity using X-ray diffraction. Diffractogram profiles of ground tissue from the *CESA8* dark zone, bright zone, and top mature zone. CI, crystallinity index.

**Fig. S6.**
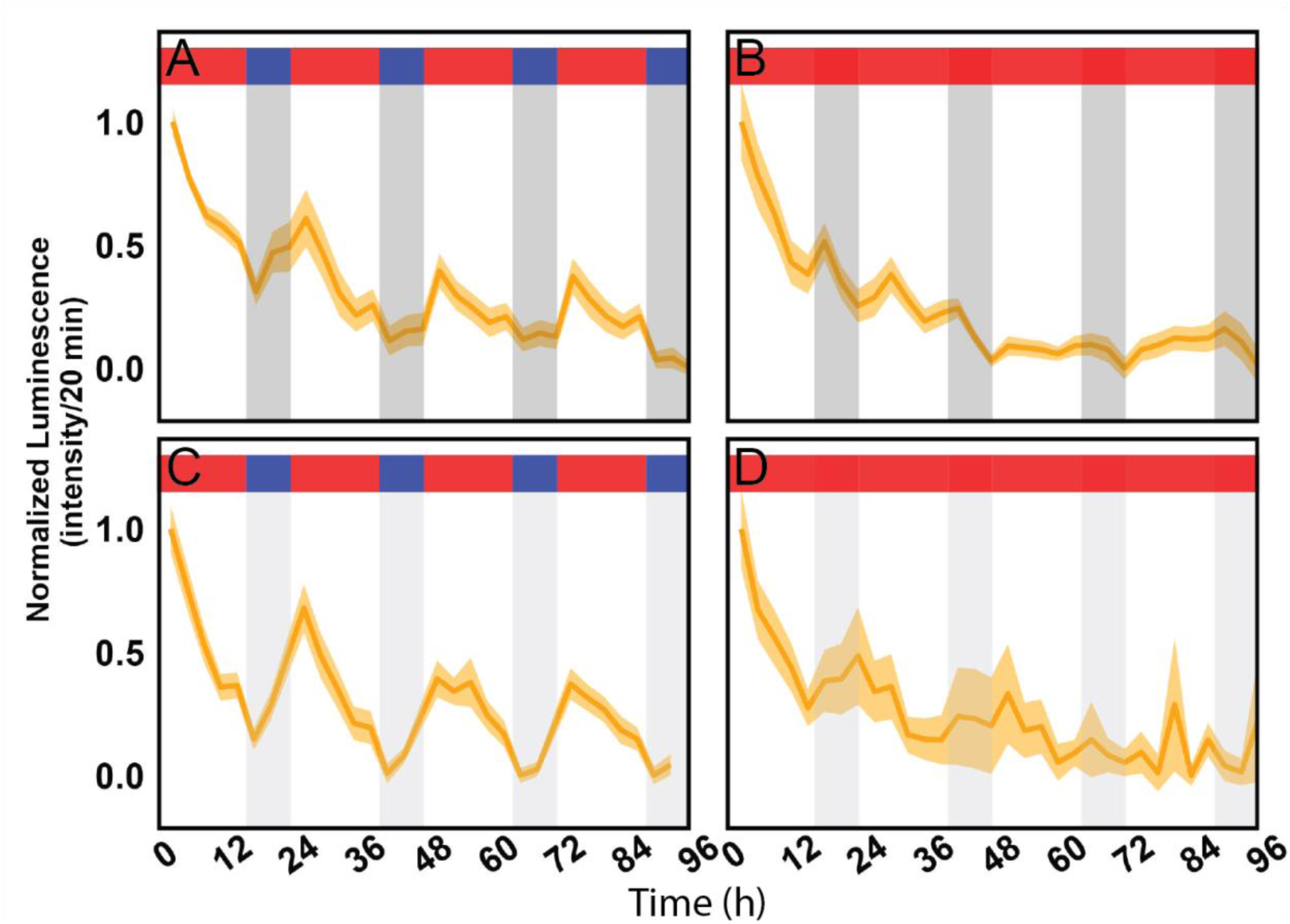
Replicate experiments of *CESA8::LUC* time course experiments. Plants were entrained under photo and thermocycles and then imaged under **(A)** photo and thermocycles, **(B)** constant light with thermocycles, **(C)** constant 25°C with photocycles or **(D)** constant light and 25°C. Luminescence data are means ± SEM, n= 7. Dark gray shading represents night, light gray shading represents subjective night, red shading indicates 25°C, and blue shading indicates 17°C. LDHC, photo and thermocycles; LDHH, photocycles and constant temperature; LLHC, thermocycles and constant light; LLHH, constant light and constant warm temperature.

**Fig. S7.**
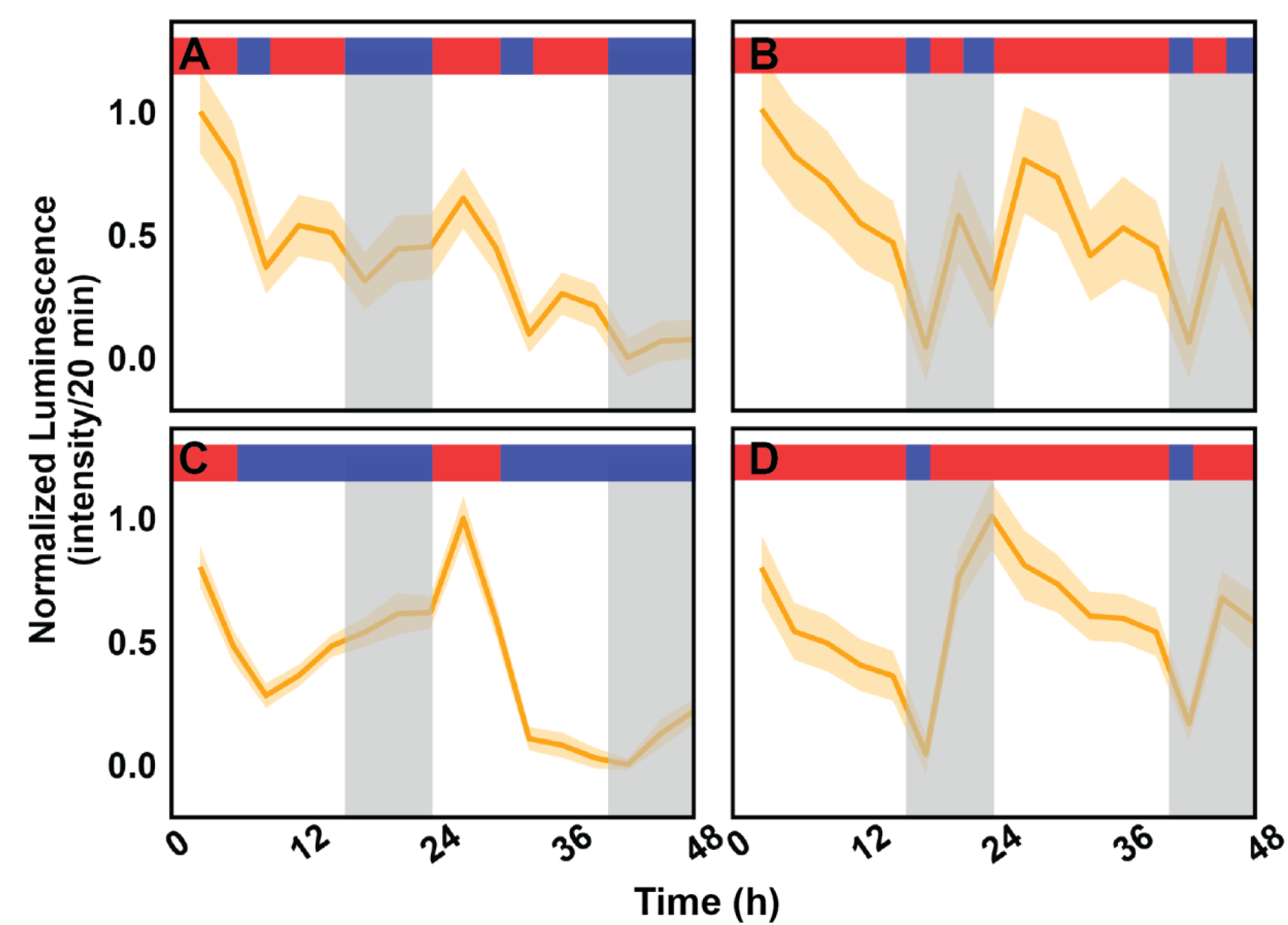
Replicate experiment of temperature pulse experiments. *CESA8::LUC* plants were entrained under photo and thermocycles, transferred to thermocycles alone and then given an (**A**) 17°C cold pulse for 3 h midday or a (**B**) 25°C warm pulse for 3 h at night. Long day entrained plants encountered (**C**) early nighttime temperature, equivalent of a short day thermocycle (6 h 25°C:18 h 17°C) or (**D**) early daytime temperature, equivalent to very long day thermocycle (21 h 25°C:3 h 17°C). Luminescence and elongation of the primary stem were measured for each timepoint. Luminescence data are means ± SEM, n= 7.

**Fig. S8.**
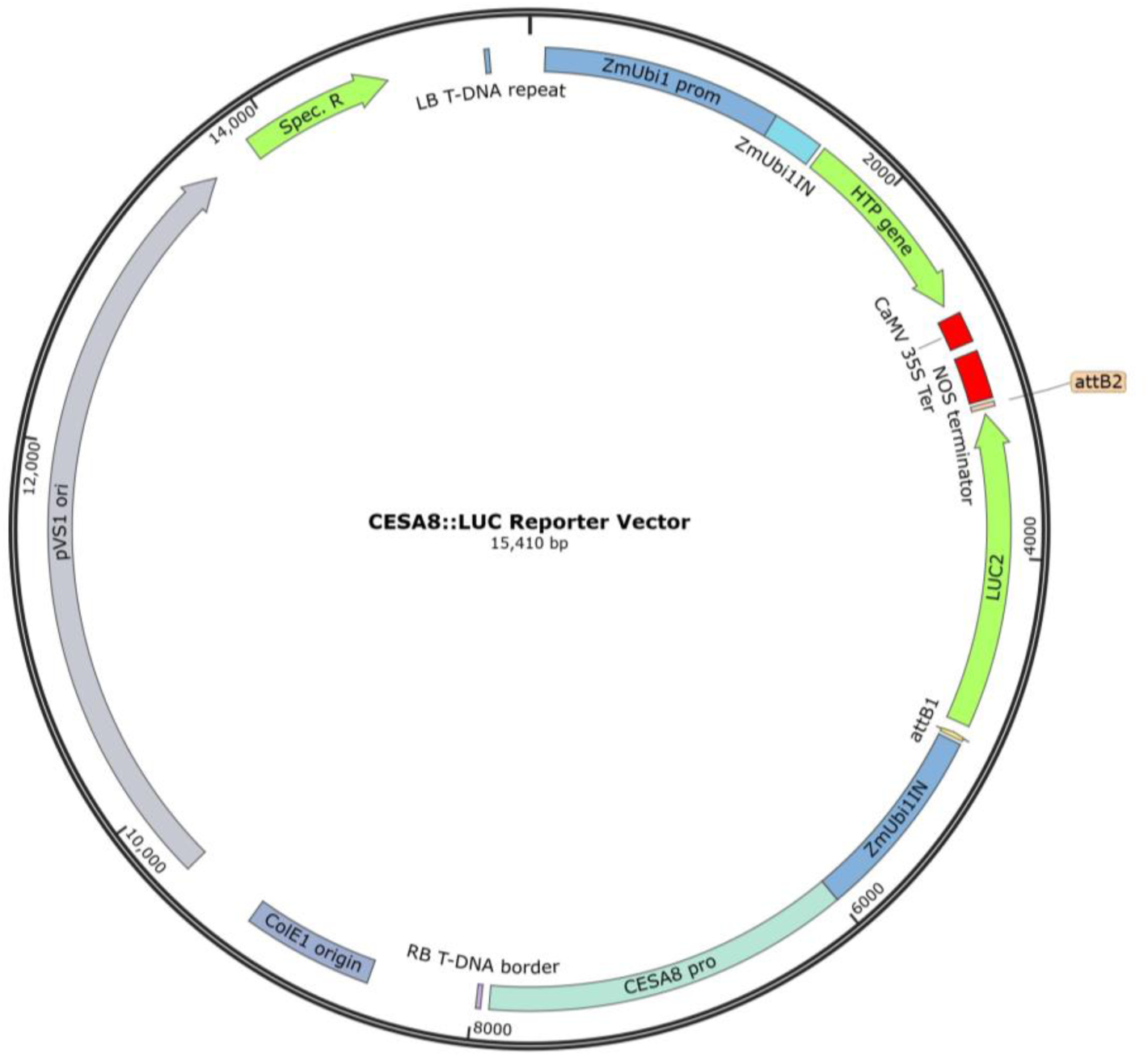
Schematic of the *CESA8::LUC* plant transformation plasmid. T-DNA, transfer DNA; LB, left border; ZmUbi prom, *Zea mays Ubiquitin1* promoter; ZmUbi1IN, *Zea mays Ubiquitin1* intron; HTP, *Hygromycin B phosphotransferase*; CaMV 35S Ter, Cauliflower mosaic virus terminator sequence; NOS, nopaline synthase; LUC, *LUCIFERASE*; attB1 and attB2, Gateway cloning sites; CESA8 pro, *CELLULOSE SYNTHASE A8* promoter; RB, right border; ColE1 origin, *Escherichia coli* origin of replication; Spec. R, spectinomycin resistance.

**Table S1.**
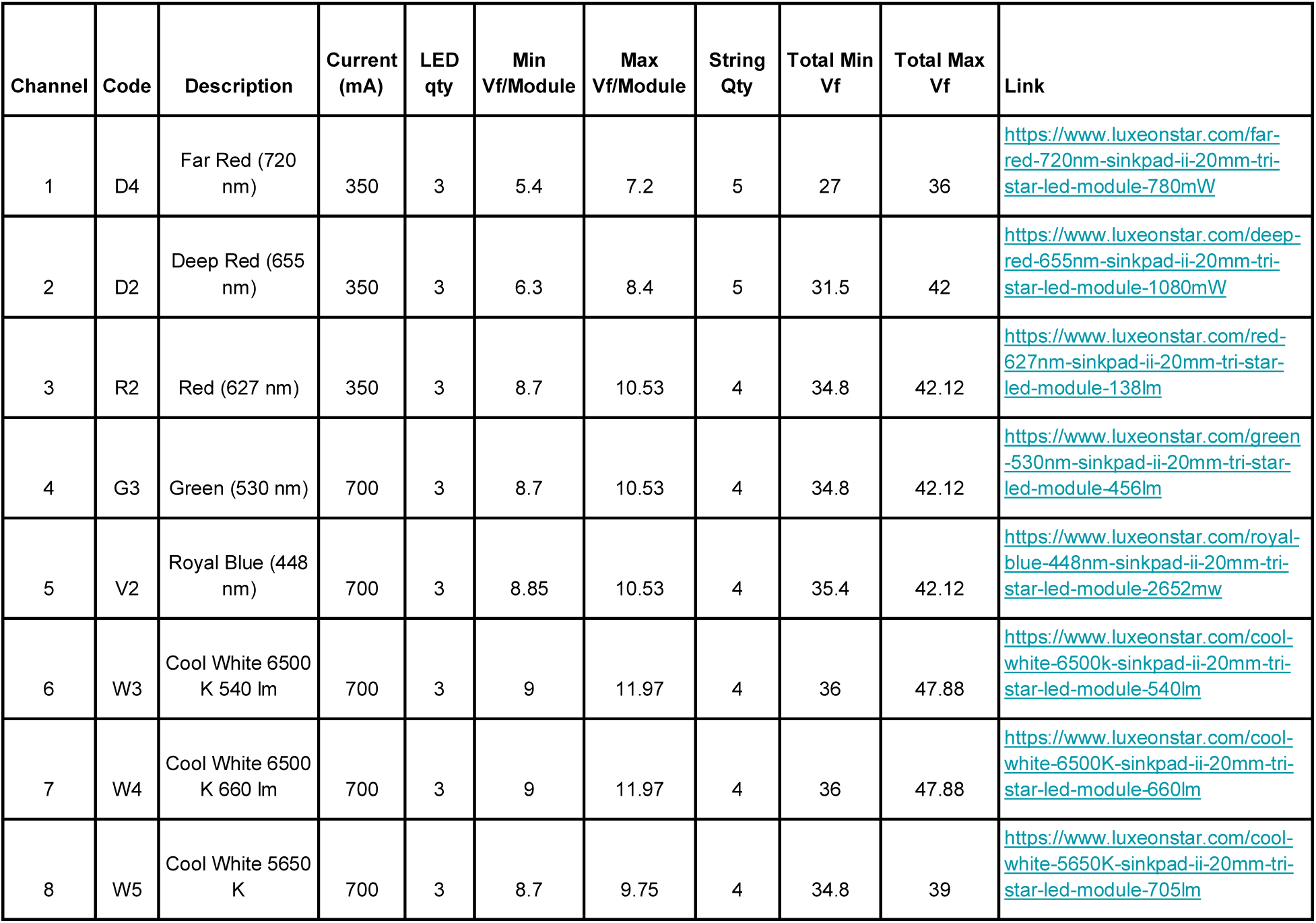
Description of light-emitting diodes in luciferase imaging chamber.

**Table S2.**
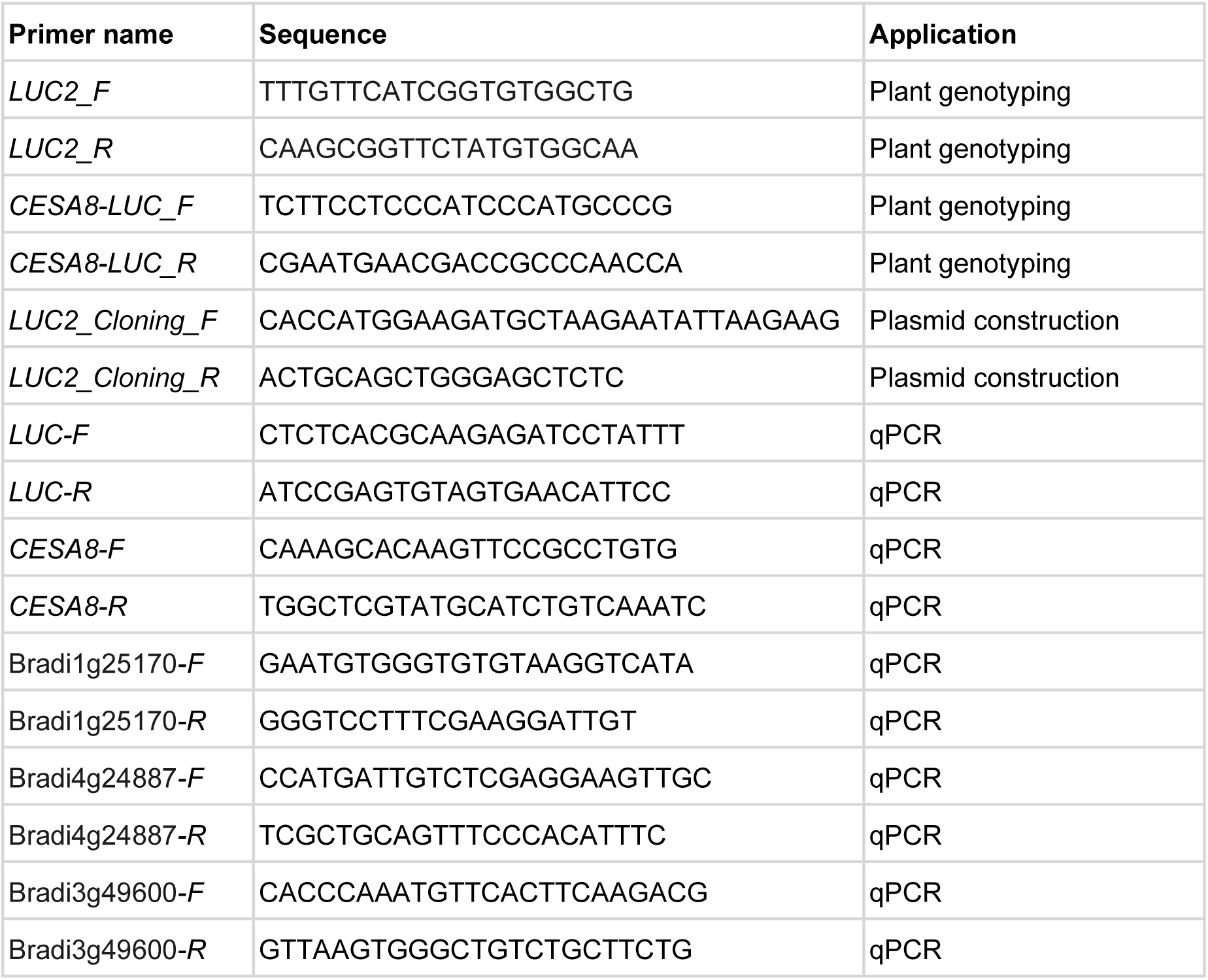
Primers used in this study.

